# Global phylogenomic analysis of *Staphylococcus pseudintermedius* reveals genomic and prophage diversity in multi-drug resistant lineages

**DOI:** 10.1101/2024.09.06.611613

**Authors:** Lucy F. Grist, Alice Brown, Noel Fitzpatrick, Giuseppina Mariano, Roberto M. La Ragione, Arnoud H. M. van Vliet, Jai W. Mehat

## Abstract

*Staphylococcus pseudintermedius* is the foremost cause of opportunistic canine skin and mucosal infections worldwide. Multidrug resistant (MDR) and methicillin-resistant *S. pseudintermedius* (MRSP) lineages have disseminated globally in the last decade and present significant treatment challenges. However, little is known regarding the factors that contribute to the success of MDR lineages. In this study, we compared the genome sequence of 110 UK isolates of *S. pseudintermedius* to 2,166 genomes of *S. pseudintermedius* populations from different continents. A novel core genome multi-locus typing scheme was generated to allow large scale, rapid and detailed analysis of *S. pseudintermedius* phylogenies, and was used to show that the *S. pseudintermedius* population structure is broadly segregated into an MDR population and a non-MDR population. MRSP lineages are predicted to either encode certain resistance genes chromosomally, or on plasmids, and this is associated with their MLST sequence type. Comparison of lineages most frequently implicated in disease, ST-45 and ST-71, with the phylogenetically related ST-496 lineage that has a comparatively low disease rate, revealed that ST-45 and ST-71 genomes encode distinct combinations of phage-defence systems and concurrently encode a high number of intact prophages. In contrast, ST-496 genomes encode a wider array of phage defence systems and lack intact, complete prophages. Additionally, we show that distinct prophages are widespread in *S. pseudintermedius* and appear to account for the vast majority of genomic diversity. These findings indicate that MRSP lineages have significant structural genomic differences, and that prophage integration, and differential antiviral systems correlate with the emergence of successful genotypes.

**Impact Statement:** *Staphylococcus pseudintermedius* is a major cause of soft tissue infections in dogs but may occasionally infect other companion animals and humans, most often veterinary professionals in primary care. Methicillin-resistant *Staphylococcus pseudintermedius* (MRSP) have emerged worldwide and are often linked to resistance to multiple antimicrobials, resulting in a significant health burden. Here, we analysed a large collection of *S. pseudintermedius* genome sequences, which has allowed a detailed characterisation of the molecular epidemiology and diversity of the species using a novel typing scheme.

Here we show that closely related MRSP lineages differ in whether specific antibiotic resistance genes are encoded on potentially mobilisable plasmids, or more stably on the chromosome, indicating differing evolutionary trajectories of MRSP lineages. Comparison of the *S. pseudintermedius* types most frequently implicated in clinical cases to those associated with commensal carriage, showed that known genes thought to contribute to disease are universal, and therefore, not associated with the high incidence rates of disease of particular lineages. We demonstrate that MRSP sequence types linked to high disease rates lack specific phage-defence systems and are associated with a high burden of prophages.

**Data statement:** All supporting data, code and protocols have been provided either within the article, through supplementary data files, or via Figshare and Zenodo. Six supplementary figures and one supplementary file are available with the online version of this article.

**Data summary:** All genome sequencing data (sequencing reads and genome assemblies) are available at the Sequence Read Archive (SRA) and Genomes repositories at NCBI, in Bioproject number PRJNA1153484. The information for the individual *S. pseudintermedius* genomes has been included in Supplementary File 1. Genome assemblies used in this study are available from https://zenodo.org/records/13692319, DOI <u>10.5281/zenodo.13692318</u>. The *S. pseudintermedius* cgMLST scheme developed for this study has been made available via Zenodo (https://zenodo.org/records/13633136, DOI: 10.5281/zenodo.13633135) and Figshare (https://doi.org/10.6084/m9.figshare.26911654.v1, DOI: 10.6084/m9.figshare.26911654).

## Introduction

*Staphylococcus pseudintermedius* is a common member of the skin and mucosal microflora of dogs and other companion animals (1). However, it is also the foremost cause of opportunistic canine infections worldwide, and is commonly implicated in pyoderma (2), otitis externa (3), post-operative bone infections and surgical abscesses (4, 5), as well as respiratory and urogenital tract infections (4, 6, 7). Consequences of infection can range from mild tissue inflammation to severe necrosis (7). Humans may play a role in transmission in reverse zoonotic transmission as 4.1% of dog owners are estimated to be carriers of this organism (8–10). Treatment of bacterial infections in companion animals often involves the same antimicrobial classes that are critical in human medicine, including first- and third-generation cephalosporins and fluoroquinolones (11). Given the close proximity and contact between humans and companion animals, there is significant potential for the evolution and transmission of antimicrobial resistance in either host.

Recent genomic studies have shown a high level of genetic diversity of *S. pseudintermedius*, with more than 1,400 multi-locus sequence types reported (12–14). However, these studies have mostly primarily focused on singular geographic regions, leaving a picture of *S. pseudintermedius* lacking diversity. Moreover, epidemiological studies tend to focus on methicillin-resistant *Staphylococcus pseudintermedius* (MRSP) isolates of clinical relevance which are often multi-drug resistant (MDR). Multidrug resistant (MDR) and methicillin-resistant *S. pseudintermedius* (MRSP) lineages have disseminated widely in the last two decades and present significant treatment challenges (2, 3, 15). However, this focus on MRSP isolates may distort understanding of *S. pseudintermedius* species diversity and consequently understanding of other factors driving evolution and emergence.

The most common sequence type implicated in *S. pseudintermedius* infections is ST-71, followed by ST-45 and ST-68 (12, 13, 16–18). Analysis of *S. pseudintermedius* populations in the USA have suggested that these MRSP genotypes arose through multiple, independent gene acquisition events-including resistance genes, followed by clonal expansion (12, 13) presumably facilitated by the use of many classes of antibiotics in veterinary medicine. MRSP isolates have been reported to encode a larger accessory genome than their MSSP counterparts (19), this suggests that gene acquisition is a primary driver in the emergence of pathogenicity.

The prevalence of genes associated with antibiotic resistance, virulence, prophages, and horizontal gene transfer has been reported to differ amongst the epidemic clones (14), and epidemiological studies clearly indicate that resistance to antimicrobials is not a pre-requisite for disease as evidenced by antibiotic-sensitive isolates implicated in disease (17). Identifying the characteristics that dictate dominance of certain genotypes in clinical cases, such as ST-71 and ST-45, over other disease-causing *S. pseudintermedius* lineages is required to understand patterns of dissemination, and improve surveillance.

In this study, we have aimed to identify the factors that contribute to the success of genotypes associated with dissemination and clinical disease, beyond the acquisition of resistance genes. To allow comparison and clustering of a large number of *S. pseudintermedius* genomes, we have generated a core-genome MLST scheme. We have used this scheme to characterise the diversity of the *S. pseudintermedius* phylogeny, and identified that MDR/MRSP lineages differ in whether specific antibiotic resistance genes are encoded on plasmids, or the chromosome. Furthermore, we show that sequence types associated with multi-continent dissemination and high rates of clinical disease encode a high density of prophages and encode a distinct combination of phage-defence systems.

## Methods

### Collection, culture and sequencing of *S. pseudintermedius* clinical isolates from the UK

We performed whole genome sequencing on 110 *S. pseudintermedius* isolates collected from a single veterinary referral practice between 2015 and 2017. All isolates were obtained from canine patients as part of routine clinical practice and confirmed as *S. pseudintermedius* by Loop-Mediated Isothermal Amplification, and culture (5). Individual isolates were cultured from cryovials stored at -80°C on to Columbia Blood Agar (Oxoid, UK), aerobically at 37°C for 24 hours. DNA was extracted using the GenElute™ Bacterial Genomic DNA Kit (Sigma, UK). Genome sequencing was provided by MicrobesNG (http://www.microbesng.com). Genomic DNA libraries were prepared using the Nextera XT Library Prep Kit (Illumina, San Diego, USA) following the manufacturer’s protocol with the following modifications: input DNA was increased 2-fold, and PCR elongation time was increased to 45s. DNA quantification and library preparation were carried out on a Hamilton Microlab STAR automated liquid handling system (Hamilton Bonaduz AG, Switzerland). Libraries were sequenced using Illumina sequencers (HiSeq) using a 250bp paired end protocol. Sequence adapters were trimmed using Trimmomatic (version 0.30) with a sliding cut-off of Q15, and assembled using Shovill version 1.1.0 (https://github.com/tseemann/shovill) using the default settings and the Spades assembler (20).

### Download of publicly available *S. pseudintermedius* genomes and associated metadata

Metadata for all available *S. pseudintermedius* genomes was downloaded from the NCBI Pathogens Detection database (https://www.ncbi.nlm.nih.gov/pathogens/). Corresponding assemblies were downloaded and quality assessed using QUAST version 4.6.3 (21). Samples for which only sequence reads were available (Supplementary File 1), were processed for assembly with Shovill/Spades as above. Only genomes with N50>50 kb, L50<20, and a number of contigs <200 were included in this study. Assemblies with an absence of information on isolation source were excluded. The final dataset of 2,276 *S. pseudintermedius* genomes (Supplementary File 1) was comprised of 110 UK genomes and 2,166 publicly available sequences representing the wider geographical population. Each assembly was categorised as disease, or non-disease, according to metadata. Samples derived from pyoderma, wounds, abscesses, or internal organs were classed as “disease”, whereas samples isolated from the skin of healthy hosts was classed as “non-disease”.

### Phylogenetic reconstruction, multi-locus sequence typing and core-genome sequence typing

The UK assemblies were typed according to the PubMLST scheme for *Staphylococcus pseudintermedius* (https://pubmlst.org/organisms/staphylococcus-pseudintermedius) using mlst version 2.23.0 (https://github.com/tseemann/mlst). Phylogenetic reconstruction of UK *S. pseudintermedius* genomes was performed based on core-genome SNPs using ParSNP version 1.7.4 (22). Due to a high degree of genotypic variation, a core-genome MLST scheme was developed using chewBBACA version 2.8.5 (23) with default settings (https://github.com/B-UMMI/chewBBACA_tutorial). A training file was generated using Prodigal version 2.6.3 (24). The scheme was generated using 74 complete genomes and is comprised of 1,356 genes present in 99% of the genomes. cgMLST types were designated relative to conventional MLST sequence types. Phylogenetic trees from the chewBBACA allele calls were constructed using GrapeTree version 1.5.0 and the RapidNJ algorithm (25). The *Staphylococcus pseudintermedius* cgMLST scheme is available from (Zonodo-DOI: 10.5281/zenodo.13633135 and Figshare-DOI: 10.6084/m9.figshare.26911654).

### Detection of virulence and antimicrobial resistance genes, and prediction of plasmid contigs

The NCBI AMRFinderPlus tool version 3.10 (26) was used in combination with Pointfinder (27) to identify antimicrobial resistance genes and resistance-associated point mutations. AMRfinder information was used to assign each genome as either MRSP (*mecA*-positive) and MSSP (*mecA*-negative). Abricate version 1.0.1 (https://github.com/tseemann/abricate) was used to search for known *S. pseudintermedius* virulence genes (17, 27) and biocide resistance genes in all genomes. Prediction of plasmid and chromosomal contigs was performed using RFplasmid (28) using the provided *Staphylococcus* analysis file. Contigs with an 60% or higher probability score were considered to have a plasmid origin.

### Pangenome-wide analysis for markers enriched in disease-associated backgrounds

All genome assemblies were annotated using Prokka version 1.14.6 (28). A pan-genome was constructed using Roary version 3.13 (30) with a BLAST cut-off of 95%. Gene markers overrepresented in genotypes associated with high clinical disease (e.g. ST-71, ST-45, ST-258), and low clinical disease (e.g. ST-496), within a narrow genetic background were identified using Scoary version 1.16 (31) with an initial threshold of a Bonferroni-corrected p-value of 0.05. Genes determined as present in >90% in the target genomes and in <10% of the non-target genomes were considered significant. Select genomes associated a high density of putative phage genes were screened using PHASTEST (32), confirming the presence of prophages. Putative phage defence systems and genes were identified using PADLOC v2.0.0 PADLOC-DB v2.0.0 using default settings.

## Results

### Population structure of *Staphylococcus pseudintermedius* isolated in the UK

Within a UK veterinary referral practice, *S. pseudintermedius* was implicated in approximately 30% of all bacterial infections arising from surgical site infections (SSIs) over a four-year period. A total of 110 isolates obtained and their genome sequence was determined, and used to analyse their population structure. Within these 110 isolates, ST-71 was the dominant sequence type accounting for around 40% of the *S. pseudintermedius* isolates in the collection. Forty-nine isolates derived from clinical *S. pseudintermedius* infections in the UK were unable to be assigned a sequence type according to the existing MLST scheme (34). These novel clonal complexes were composed of highly diverse genotypes indicative of a highly varied population structure of disease-causing isolates in the UK, beyond the dominant ST-71 lineage (Figure 1).

**Figure 1:**
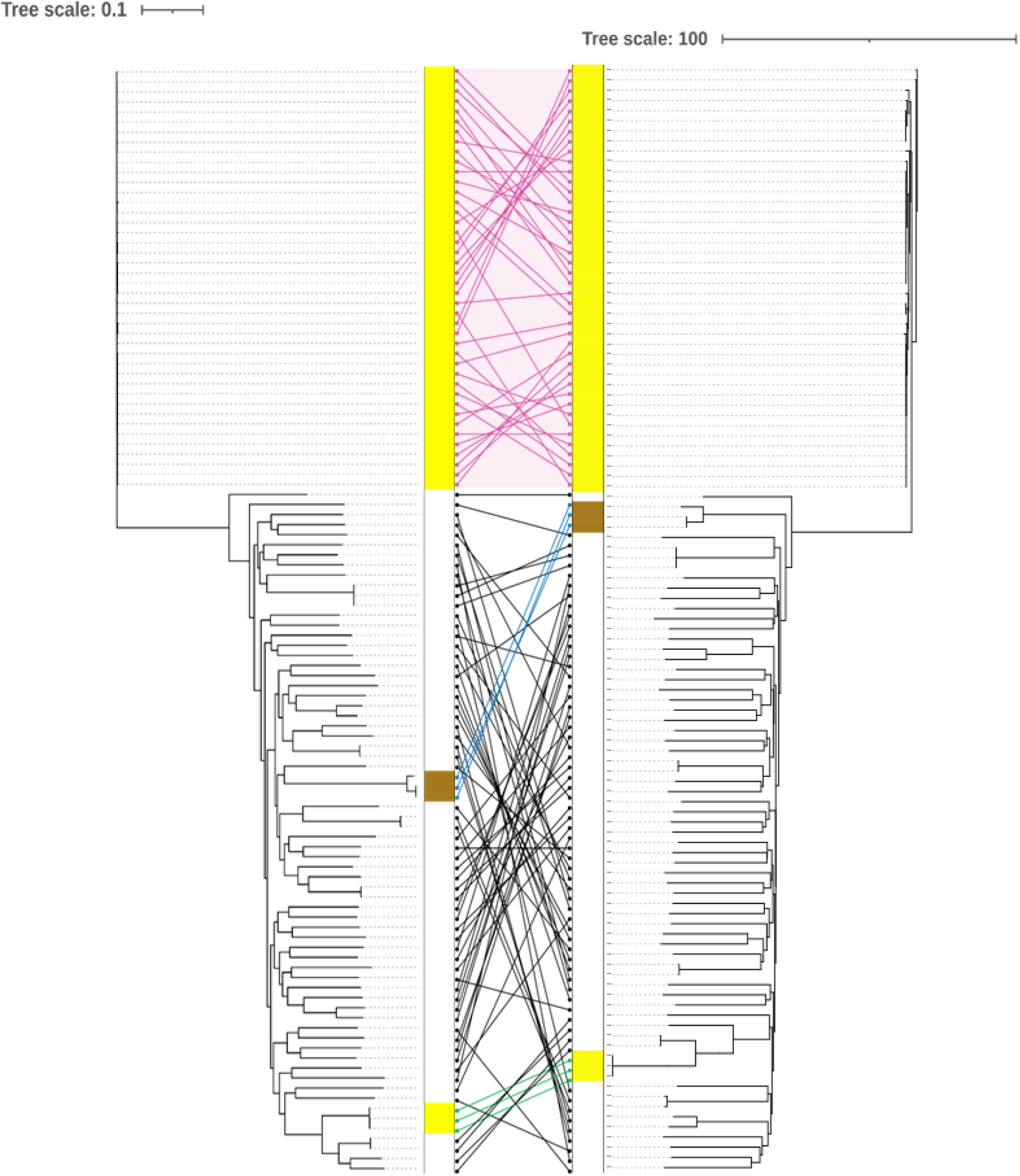
Tanglegram showing a comparison of phylogenetic reconstruction of 110 *S. pseudintermedius* genomes collected in this study from clinical cases in the UK as part of routine clinical practice. The tanglegram shows a comparison of core-genome SNPs using ParSNP (left), and a reconstruction based on cgMLST using Grapetree (right). Phylogenetic analysis based on the cgMLST scheme correlated well with the existing pubMLST scheme, and clustering of major clonal complexes such as ST-71 (pink shading and connecting lines), is largely conserved.

A total of 57.2% of our UK clinical isolates were classified as multi-drug resistant (MDR) and 51% were determined to be MRSP. Beyond ST-71, the MDR/MRSP trait was also found in disparate MLST types including ST-45, ST-277 and ST-301. To more accurately characterise and interrogate the dataset, a core-genome MLST scheme was generated and implemented which allowed discriminative typing of all genomes. Phylogenetic analysis based on the cgMLST scheme correlated well with the existing pubMLST scheme and phylogenetic typing based on core-genome SNPs (Figure 1), and as cgMLST allows for analysis of larger genome data sets, all subsequent phylogenetic analyses were based on cgMLST.

### Pan-genome analysis of Staphylococcus pseudintermedius

An additional 2,166 *S. pseudintermedius* genomes acquired from the NCBI Pathogens Detection database were acquired to supplement the UK *S. pseudintermedius* dataset and construct a pan-genome more representative of *S. pseudintermedius* genomes circulating world-wide (Supplementary File 1). Virulence gene and antimicrobial resistance gene distribution within the 2,276 *S. pseudintermedius* genomes was determined within the context of associated metadata for each genome. Genomes were classified as multi-drug resistant if intact resistance genes to 3 or more different classes of antibiotics were detected. The major MDR genotypes were determined and correlation of MDR with disease and geographical site of isolation was evaluated.

Of 8,809 unique genes within the pan-genome, only 1,995 were classified as core genes. Remarkably, of the 6,814 accessory genes identified, 6,099 were only present 0-15% of genomes. This suggests that the majority of accessory genes are lineage-specific, or even unique to single isolates, and not shared amongst all genotypes despite occupying the same niche. Consistent with this, the rate at which novel accessory genes were detected remained constant as genomes were incorporated into the pan-genome (Figure S1), indicating that *S. pseudintermedius* has an open pan-genome characterised by high phylogenetic diversity and acquisition of novel genes through horizontal transfer. This suggests that acquisition of novel genetic material is widespread in the species and may profoundly impact population structure.

### Epidemiology, clinical disease association, and multi-drug resistance of ***Staphylococcus pseudintermedius* genotypes**

The 2,276 *S. pseudintermedius* genomes are comprised of a range of commensal and disease-causing isolates. However, the dataset is heavily skewed towards genomes derived from the USA; consequently, there are many genotypes that appear to be geographically restricted, particularly within the MDR cluster, for example ST-155, ST-64 and ST-551. In contrast, there are clonal complexes that are widely disseminated, including ST-71, ST-45 and ST-258, and to a lesser extent ST-496, which are also MDR lineages (Figure 2).

**Figure 2:**
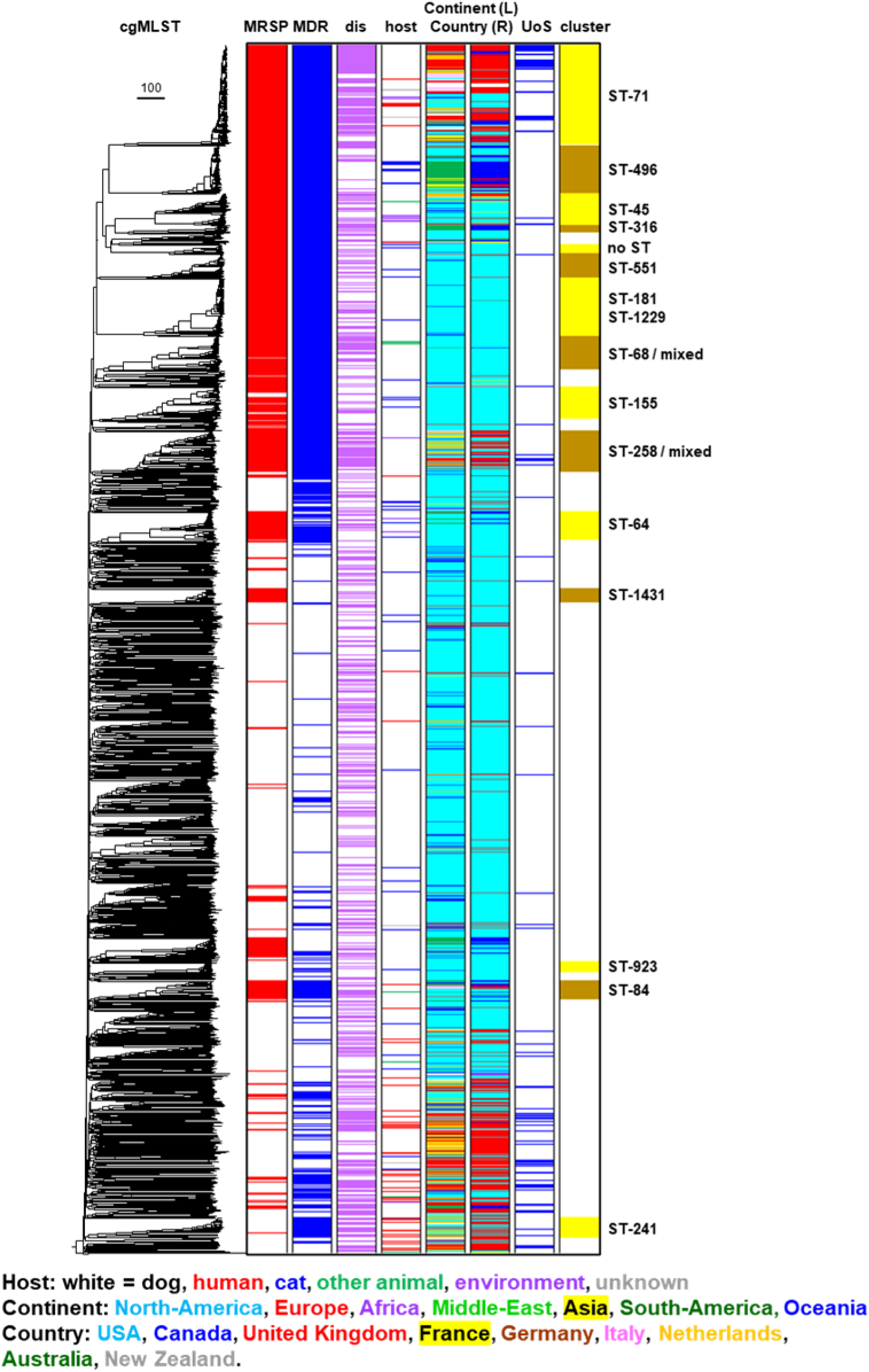
cgMLST phylogeny of 2276 *Staphylococcus pseudintermedius* assembled genomes (Supplementary File 1) comprised of a range of commensal and disease-causing isolates (purple shading, column labelled “dis”), predominantly originally isolated from dogs (white shading, column labelled “host”. The dataset encompasses genomes derived from multiple continents; North-America, Europe, Africa, Middle-East, Asia, South-America, Oceania. Genomes sequenced as part of this study are coloured blue in the UoS column, and publicly available genomes from sequence repositories are white in this column. Genomes of isolates classed multi-drug resistant on the basis of encoding genes conferring resistance against 3, or more, classes of antibiotics are shaded blue in column labelled “MDR”. Genomes classes as MRSP are coloured red in the “MRSP” column. The MDR/MRSP phylogenetic clusters are predominantly composed of ST-71, ST-496, ST-45, ST-316, ST-551, ST-181, ST-1229, ST-68, ST-155, and ST-258.

Almost one-third (32%) of the *S. pseudintermedius* genomes in the whole dataset are associated with clinical disease (Table 1A). However, certain clonal complexes were over-represented in this disease-associated group (Figure 3; Table 1B). More than two-thirds (70%) of ST-71 genomes are associated with clinical disease. Conversely, ST-496, which is phylogenetically closest to ST-71, consists of genomes that are primarily derived from non-diseased hosts (86%), indicating that this clonal complex is strongly associated with commensal carriage.

**Table 1:** The proportion of 2274 *S. pseudintermedius* genomes used in this study that are classified as either MRSP or MSSP, and their association with commensal carriage (non-disease) or clinical (disease) cases.

| MRSP/MSSP | Non-disease | Disease | Total |
| --- | --- | --- | --- |
| MRSP | 569 | 389 | 958 (42%) |
| MSSP | 978 | 340 | 1318 (58%) |
| **Total** | **1547 (68%)** | **729 (32%)** | **2276** |

**Table 2:**
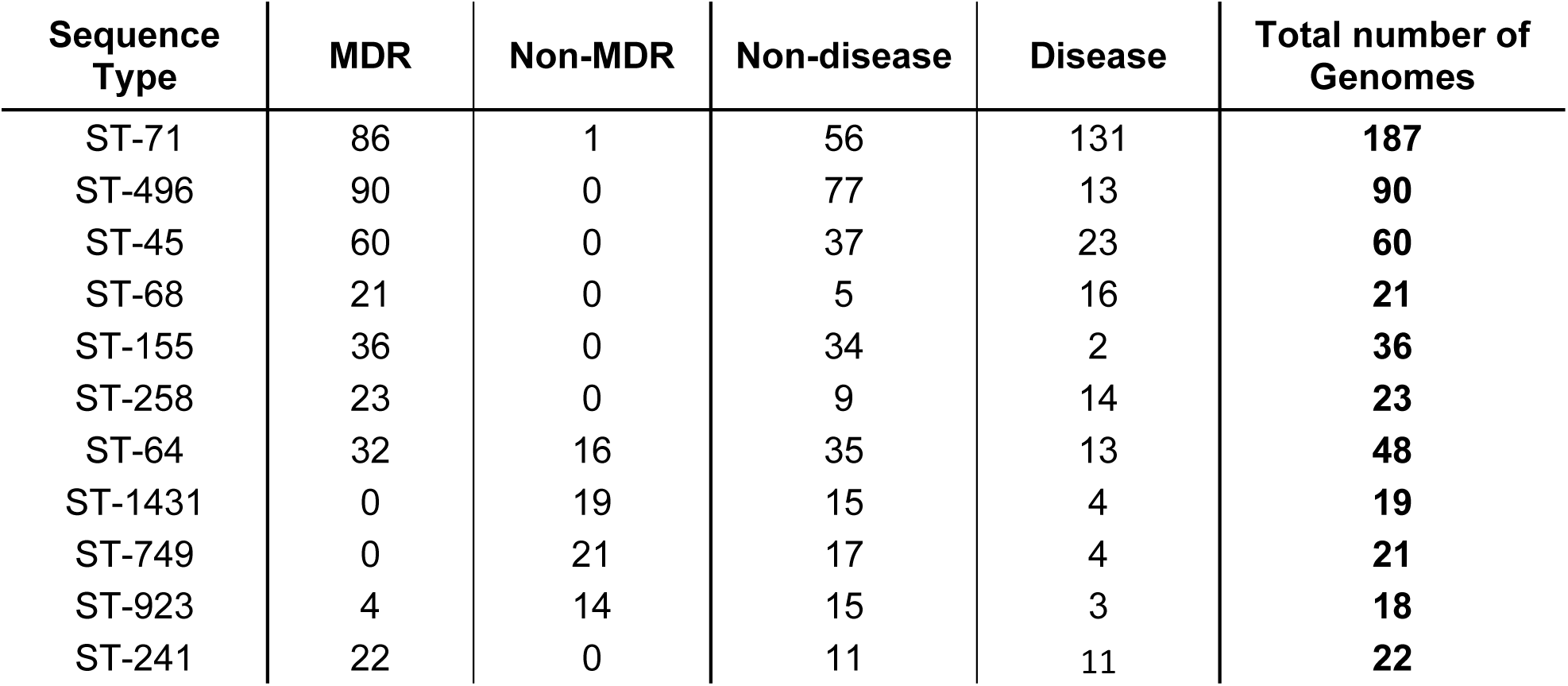
A breakdown of prevalent *S. pseudintermedius* sequence types and their association with multi-drug(MDR) resistant status, and rates of clinical cases (disease).

| Sequence Type | MDR | Non-MDR | Non-disease | Disease | Total number of Genomes |
| --- | --- | --- | --- | --- | --- |
| ST-71 | 86 | 1 | 56 | 131 | **187** |
| ST-496 | 90 | 0 | 77 | 13 | **90** |
| ST-45 | 60 | 0 | 37 | 23 | **60** |
| ST-68 | 21 | 0 | 5 | 16 | **21** |
| ST-155 | 36 | 0 | 34 | 2 | **36** |
| ST-258 | 23 | 0 | 9 | 14 | **23** |
| ST-64 | 32 | 16 | 35 | 13 | **48** |
| ST-1431 | 0 | 19 | 15 | 4 | **19** |
| ST-749 | 0 | 21 | 17 | 4 | **21** |
| ST-923 | 4 | 14 | 15 | 3 | **18** |
| ST-241 | 22 | 0 | 11 | 11 | **22** |

**Figure 3:**
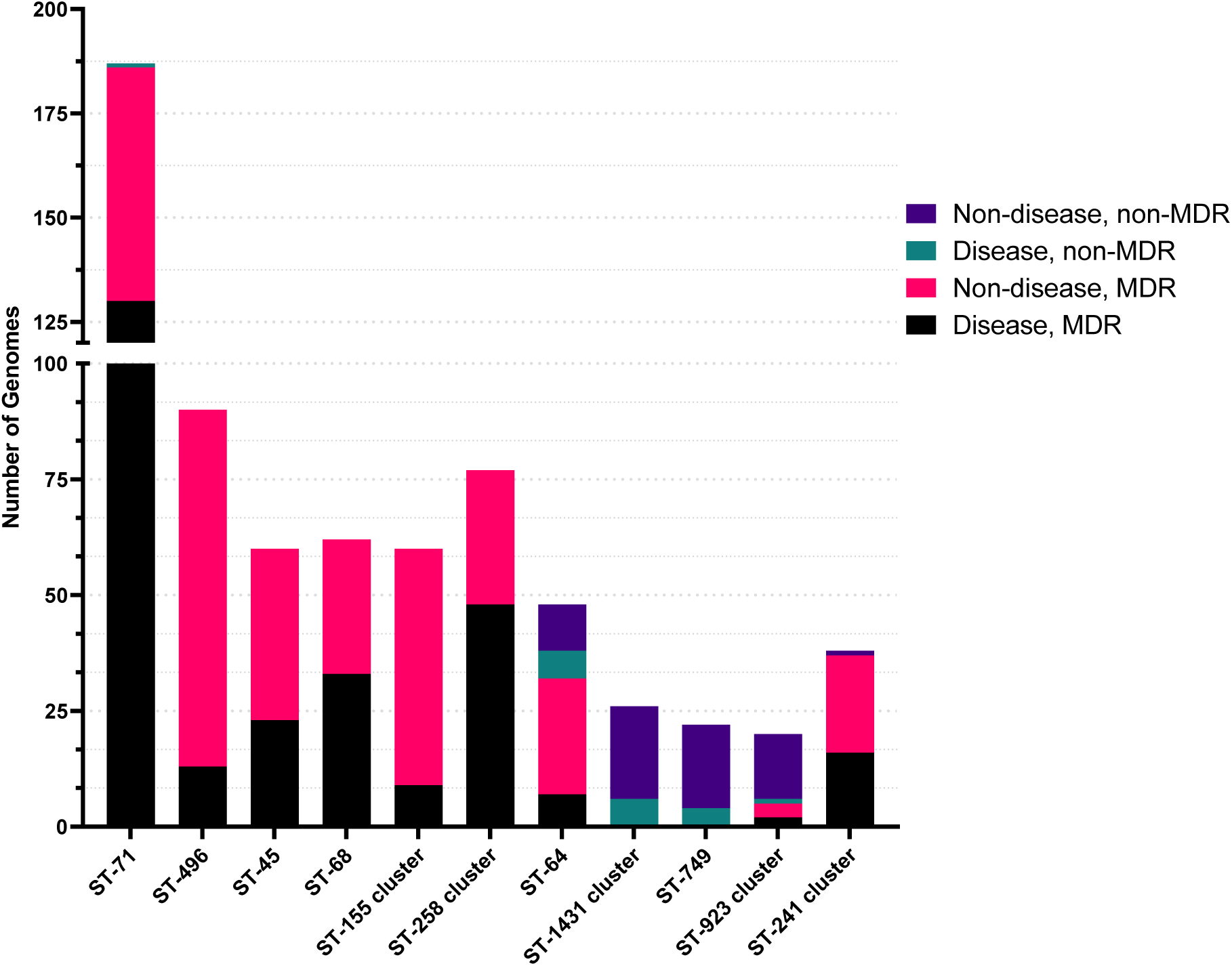
Stacked bar plot showing *S. pseudintermedius* sequence type association with disease and multi-drug resistant status. ST-71 is an MDR lineage that is strongly associated with disease. Conversely, ST-496 is a closely related MDR sequence type that is associated with commensal carriage. ST-1431, ST-749, and ST923 are non MDR lineages that are more commonly associated with commensal carriage.

Of the genomes derived from clinical samples, 51% (424/729) are classified as MDR. Of the non-clinical samples, 45% (n=699/1547) were classified as MDR. There appears to be no combination of resistance genes that is more associated with either disease or non-disease genomes. This shows that MDR-status is not a pathogenic trait in itself, and is not a predictor of clinical disease.

The *S. pseudintermedius* phylogeny can be broadly segregated into an MDR population and a non-MDR population. Each contig from the 2276 assemblies in this collection was classified as either plasmidic or chromosomal based on the Rfplasmid score; this revealed that the MDR population varies significantly in how resistance genes are encoded (Figure 4). Beta-lactamase resistance is encoded across the phylogeny, most often on the chromosome, but also on both the chromosome and a plasmid, as in the case of a ST-496 sub-population. Aminoglycoside resistance is associated with the MDR population and is encoded by plasmids or both a plasmid and on the chromosome. Tetracycline resistance is chromosomally encoded across the phylogeny except in ST-71 where it is encoded on a plasmid. ST-496 and ST-551 which encode tetracycline resistance on both the chromosome and on a plasmid. The ST-496, ST-45, ST-316, and ST-68 clusters encoded macrolide resistance on predicted plasmid contigs, whereas most other MDR lineages are predicted to encode macrolide resistance on the chromosome. Similarly, trimethoprim resistance is encoded chromosomally by MDR lineages with the exception of ST-68, which encodes this resistance on predicted plasmid contigs.

**Figure 4:**
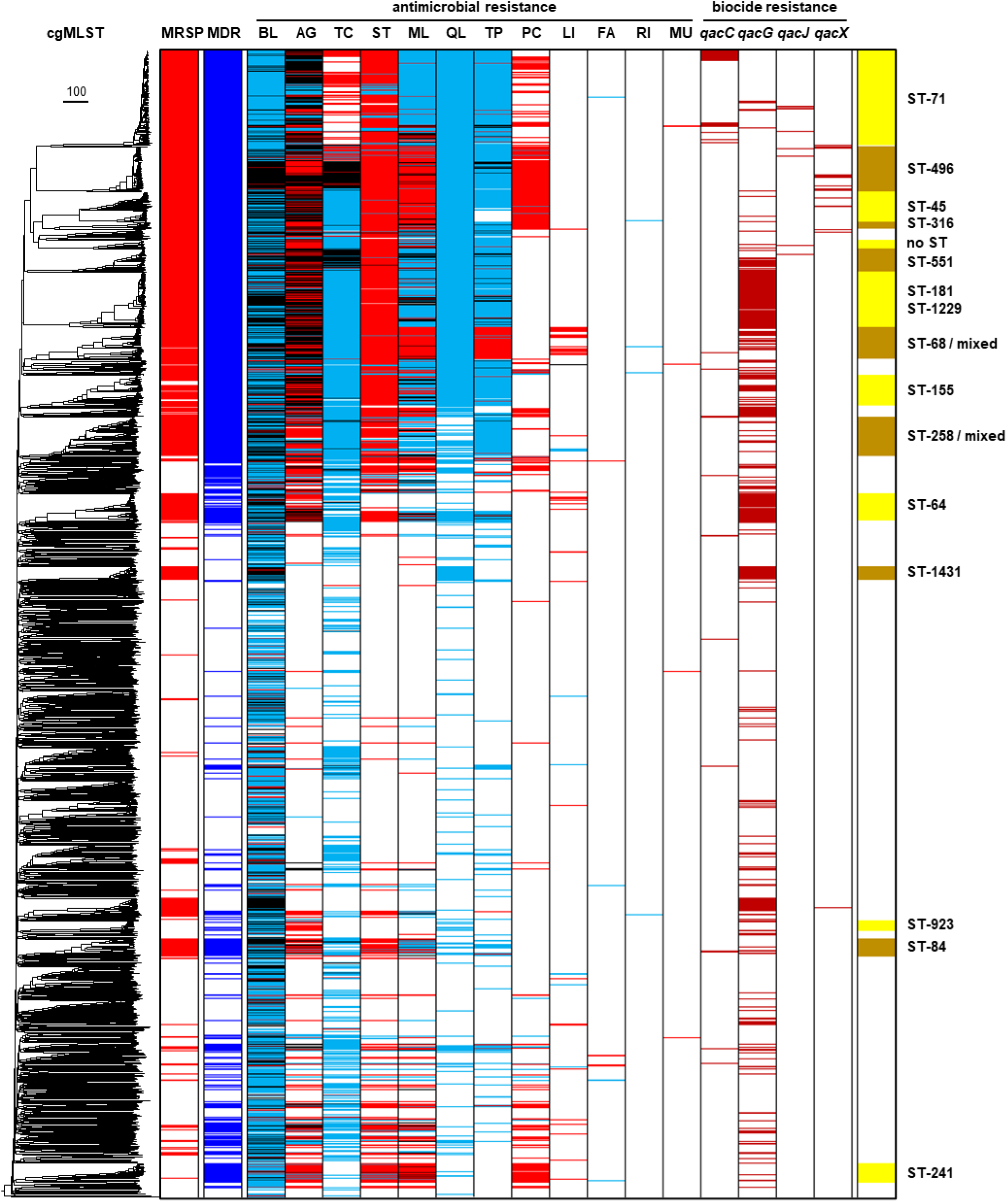
The phylogenetic distribution of antimicrobial resistance (AG = aminoglycoside, BL = beta-lactam, FA = fusidic acid, LI = lincosamide, ML = macrolide, MU = mupirocin, PC = phenicol, QL = quinolone, RI = rifampicin, ST = streptothricin, TC = tetracycline, TP= trimethoprim), and biocide (*qacC, qacG, qacJ, qacX*) genes in *Staphylococcus pseudintermedius*. Each gene was predicted to be encoded on either the chromosome (blue shading), plasmid (red shading), or both (black shading). Sequence types differ in the manner by which resistance genes are encoded; tetracycline resistance is chromosomally encoded across the phylogeny except in ST-71 where it is encoded on a plasmid. Trimethoprim resistance is encoded chromosomally by MDR lineages with the exception of ST-68. The ST-496, ST-45, ST-316, and ST-68 clusters encoded macrolide resistance on predicted plasmid contigs, whereas most other MDR lineages are predicted to encode macrolide resistance on the chromosome. Full details of the phylogenetic distribution of antimicrobial resistance and biocide genes are provided in Supplementary File 1.

Virulence gene content showed no association with disease genomes and was not different between major lineages (Figure S2) suggesting all genotypes are capable of causing opportunistic disease via intrinsic mechanisms encoded across the phylogeny. Virulence gene density therefore cannot account for the dominance of ST-71, ST-68, and ST-258 relative to other genotypes. In light of this, we sought to identify genetic factors that contribute to the success of ST-71 and ST-45 beyond an MDR/MRSP status.

### Prophages are major determinants of *S. pseudintermedius* diversity and reflect the distribution of phage defence systems

Genome-wide association studies were performed to identify genetic markers that are enriched in phylogenomic clusters within the MDR phylogenetic cluster associated with multiple country incidence and a high degree of clinical disease, such as the genotypes ST-71, ST-45, and ST-496 (Figure 2). Concurrently, we used this approach to identify markers associated with genotypes with a comparatively lower rate of clinical disease. We determined the distribution of these markers across the phylogeny, and assessed their impact on population structure. Many of the genes over-represented in ST-71 and ST-45 were annotated as phage genes. We used PHASTEST to characterize these putative prophages (Figure S4, S5, S6).

Large prophage sequences were primarily associated with ST-71, ST-45 and ST-258, the lineages most frequently implicated in clinical disease (Figure 5). One of these prophages, designated here as SPpB, is 28.1Kb in length and is chromosomally integrated in ST-71, and in ST-45 adjacent to core genes involved in cell division (*whiA*) and metabolism (Figure S4). There is a large (2Kb) intergenic space upstream of the SPpB integrase and upstream of the start codon on the antisense strand, so it is unlikely that SPpB integration at this site has influenced transcription of adjacent genes.

**Figure 5:**
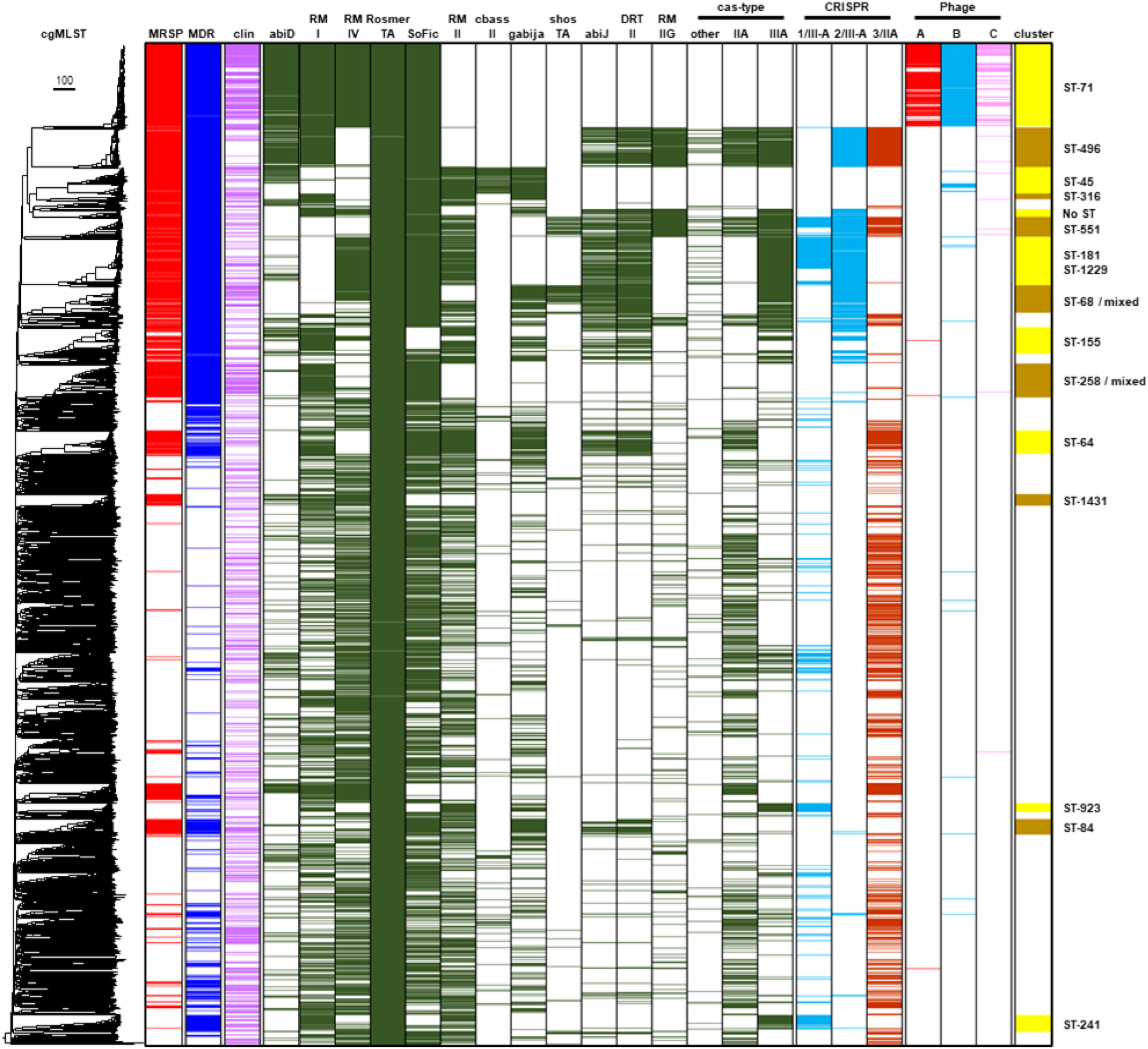
The combination of phage defence systems encoded by *S. pseudintermedius* reflects the density of prophages detected in the genomes. All 2276 genomes were screened for presence of specific prophages detected by PHASTEST, and PADLOC was used to detect antiviral genes and systems (abiD = Abortive infection system protein D, RM I = Type I restriction-modification system, RM IV = Type IV restriction-modification system, RosmerTA = RosmerTA system encoded by *rmrT* WP_231741552.1 and *rmrA* WP_058719324.1, SoFIC = SoFIC antiviral gene WP_242883646.1, RM II = Type 2 restriction-modification system, cbass II = CBASS operon composed of 4 genes, Gabija = Gabija phage defence system composed of GajA and GajB, shoshTA = shoshTA toxin/antitoxin system, abiJ = Abortive infection system protein J, DRT II = Type 2 Defence-associated Reverse Transcriptase, RM IIG = Type 2G restriction-modification system, cas-type/CRISPR = CRISPR-Cas systems). ST-71 encodes SpST71A, SPpB, and SPpC prophages (columns labelled Phage A-red colouring, B-blue colouring, and C –pink colouring), which are largely absent from other lineages. ST-71 also encodes a distinct combination of phage defence genes (indicated by green colouring) relative to most other lineages. ST-496 which has a dearth of prophage sequences, encodes a unique combination of antiviral genes. Full details of the phylogenetic distribution of identified phage defence systems and prophages are provided in Supplementary File 1.

An additional 43.7kb complete prophage associated with certain MRSP backgrounds was identified. This prophage, designated here as SPpC, appears to have integrated in the genomes of *S. pseudintermedius* ST-71, but exhibits variation in ST-45 and ST-68 and is significantly truncated in other genetic backgrounds (Figure 5). SPpC is inserted adjacent to core genes, including the *suf* operon involved in oxidative stress repair, and *corB*, involved in Mg^2+^ and Co^2+^ transport, as well as tRNA genes which are common targets for mobile genetic elements in bacterial genomes (35).

In contrast to related clonal complexes such as ST-71 and ST-45, intact prophages were not detected in ST-496. Seeking explanation for the lack of intact prophages in this lineage, GWAS determined that ST-496 encodes two independent Type III-A CRISPR-Cas systems absent from ST-71 and ST-45. Following this, we used PADLOC to systematically detect known anti-phage systems across the phylogeny, and found that the repertoire of phage-defence systems encoded differs markedly between these closely related lineages. ST-71 and ST-45 encode fewer phage defence systems than any other lineage (Supplementary File 1). Furthermore, the combination of the antiviral genes they do possess are distinct relative to other MDR-associated sequence types. ST-71 encodes the abortive infection system AbiS, two restriction-modification (RM) systems (type I and IV), and SoFic, in addition to the RosmerTA defence system encoded by all *S. pseudintermedius* (Figure 5). ST-496 lacks the Type IV RM system, but also encodes type-IIA and type-IIIA CRISPR-Cas systems, AbiJ, DRT class II, and type IIG RM system. ST-45, on the other hand predominantly encode a cbass type IIs, and type II RM system, and Gabija, as well as the occasional instance of either AbiD, or a type I RM system. The variation in the number and type of phage defence systems encoded by these related lineages reflects the presence and absence of SpST71A, SPpB, and SPpC, and is consistent with a scenario wherein the combination of defence systems encoded by ST-496 confers stronger resistance to prophage insertion.

## Discussion

In this study, we present a comprehensive population genomic analysis of *S. pseudintermedius* using a novel core-genome multi-locus sequence typing scheme, which has facilitated the classification of novel genotypes. The *S. pseudintermedius* phylogeny is broadly segregated into an MDR/MRSP population and a non-MDR/MRSP population, indicating that despite shared niche access to a variety of mobile antimicrobial resistance genes, carriage of certain resistance genes appears to be restricted to select lineages. Within the MDR/MRSP population, we have identified MRSP lineages that exhibit differences in the means by which select resistance genes are encoded, either chromosomally or on plasmids. This indicates a degree of genomic variation within this population that is linked to genetic background.

Consistent with previous reports (14, 15, 17), our results indicate co-circulation of a diverse range of lineages and an open pan-genome consistent with accumulation of genetic material via horizontal gene transfer (HGT). One important mechanism of HGT is prophage acquisition, wherein phage genomes integrate into the bacterial chromosome. Our results suggest that prophage integration between otherwise closely related MRSP lineages accounts for most of the dissimilarity, and the presence of particular prophages in ST-71, ST-45 and ST-258 is correlated with higher disease rates. Within this MDR population, the absence of intact prophages in ST-496, which is associated with commensal carriage, is likely due to a unique array of phage-defence systems which may have rendered ST-496 resistant to phage infection and lysogenic carriage of SpST71A (14), SPpB, and SPpC. Considering prophages represent the major genome divergences between these lineages, the differences in the antiviral arsenal encoded by specific lineages appears to have had significant impact on the evolution of MRSP.

Aside from contributing to genetic diversity, bacteriophages often impact phenotypically on their host as observed in *Staphylococcus aureus* wherein bacteriophages and other mobile genetic elements mediate the transfer of pathogenicity islands, conferring new phenotypic traits that enable bacterial adaption (36). For example, the Sa3int phages, which are the most prevalent of *S. aureus* phages, carry the immune evasion cluster which encode immunomodulatory proteins Sak, Scin, and CHIPS, which act in concert to enable within-host survival (37, 38).

Prophage carriage is also associated with bacterial virulence in many other species; for example, scarlet fever is caused by specific strains of *Streptococcus pyogenes* that encode a phage derived toxin; only *S. pyogenes* that are lysogenic for these phages are able to cause scarlet fever (39). Similarly, in *Corynebacterium diptheriae,* its main virulence factor, the diphtheria toxin, is encoded on a prophage (40)*. Escherichia coli* O157:H7 and O104:H4 have acquired the Shiga-toxin encoding gene *stx2a,* through lysogeny (41). Investigating the impact of prophage sequences identified in this study on *S. pseudintermedius* pathobiology is an attractive future direction for *S. pseudintermedius* studies.

The large prophages, SPpB and SPpC, which have integrated into ST-71, ST-45 and ST-68 isolates, may contribute to the high disease-rate associated with these genetic backgrounds, relative to ST-496 which does not encode these prophages. Whilst *S. pseudintermedius* prophages themselves may harbour, and co-select for, genes that contribute to the high rate of disease caused by ST-71 and ST-258, the site of phage integration in the genome may also play a role. Integration of prophages into core-gene regions has been shown to have an impact on expression levels of adjacent genes as observed previously (42–45) and can consequently be expected to contribute to physiological changes. This is exemplified by the SpST71A prophage unique to ST-71 which is inserted into the *comG* gene and is suspected to disrupt natural genetic competence of this lineage (14). The quiescent prophages identified in this study do not appear to have interrupted any identifiable gene but are adjacent to core-genes involved in cell division, oxidative stress repair, and metal transport. The integration and expression of prophages is a complex series of interactions which involved pirating host cell transcriptional machinery (46), which may impact downstream and upstream genes.

Ultimately, it is clear that multiple phage integration events have affected genome architecture in the different lineages, which can affect phenotypic variation. The maintenance of multiple intact prophages in ST-71, 45 and 258 genomes comparative to closely related sequence types suggests these may confer a functional advantage to this lineage. Future work should aim to decipher the impact of these bacteriophages on the *S. pseudintermedius* lineages that encode them, in the context of pathogenicity and evolution.

We have observed remarkable differences between closely related genotypes in the replicon type-chromosome or plasmids-of specific genes conferring resistance to macrolides, trimethoprim, and tetracycline, highlighting differences in genomic architecture. Plasmids encoding resistance determinants can be expected to be favoured by bacterial cells over chromosomally-encoded resistance genes, if the costs of maintenance and expression are lower. Plasmid mediated resistance genes have a higher proclivity for intraspecies and interspecies transfer, which has been observed in *Staphylococci* (47), than those encoded on chromosomes. Results presented here indicate that the capability to encode a macrolide, trimethoprim, or tetracycline resistance gene on either the chromosome, plasmid, or both, is linked to sequence type and that these predicted plasmids are restricted to permissible genetic backgrounds. The sequence types that encode resistance genes on plasmids are likely to be responsible for lateral, interspecies transfer of antibiotic resistance genes.

Given these resistance genes are predicted to be plasmid-encoded, they may be present at a higher copy-number than their chromosomal counterparts in other *S. pseudintermedius* lineages, and thereby confer higher levels of resistance to antibiotics used in treatment of canine pyoderma and surgical-site infections. Distribution and genetic association of plasmid-encoded resistance genes is therefore likely to impact on treatment efficacy.

This study has identified that mobile genetic elements such as prophages and plasmids are the primary determinants of diversification within *S. pseudintermedius*, and these indicate differing evolutionary trajectories of various lineages. Considering the association of prophages with lineages most frequently implicated in disease, a greater understanding of the impact of prophage genes and integration sites on *Staphylococcus pseudintermedius* biology is required.

## Acknowledgements

We gratefully acknowledge all of the veterinary practitioners who were involved in collection of bacterial samples.

## Ethical Approval

Collection of clinical samples was performed as part of routine clinical practice, and identification and banking of bacterial isolates was approved by the University of Surrey’s animal ethics committee (NERA-2017-009-SVM)

## Conflict of Interest

The authors declare that there are no conflicts of interest.

## Funding information

This work received no specific grant from any funding agency, but was supported by the University of Surrey and Fitzpatrick Referrals. The authors disclose that Professor Noel Fitzpatrick is Director and Clinical Chair of Fitzpatrick Referrals.

## Author contributions

LG, AvV, and JM designed the study. AB, LG, and JM carried out DNA extraction. LG, GM, AvV, and JM carried out analyses and interpreted data. JM drafted the manuscript. All authors critically reviewed and approved the final version of the manuscript.

## Notes

https://zenodo.org/records/13692319

https://zenodo.org/records/13633136

### Competing Interest Statement

The authors have declared no competing interest.

## References

1 Bannoehr J, Guardabassi L. *Staphylococcus pseudintermedius* in the dog: taxonomy, diagnostics, ecology, epidemiology and pathogenicity. Vet Dermatol. 2012;23(4):253–66, e51-2 DOI:10.1111/j.1365-3164.2012.01046.x.

2 Wang Y, Yang J, Logue CM, Liu K, Cao X, Zhang W, Shen J, Wu C. Methicillin-resistant *Staphylococcus pseudintermedius* isolated from canine pyoderma in North China. J Appl Microbiol. 2012;112(4):623–30 DOI:10.1111/j.1365-2672.2012.05233.x.

3 Dziva F, Wint C, Auguste T, Heeraman C, Dacon C, Yu P, Koma LM. First identification of methicillin-resistant Staphylococcus pseudintermedius strains among coagulase-positive staphylococci isolated from dogs with otitis externa in Trinidad, West Indies. Infect Ecol Epidemiol. 2015;5:29170 DOI:10.3402/iee.v5.29170.

4 Diribe O, Thomas S, AbuOun M, Fitzpatrick N, La Ragione R. Genotypic relatedness and characterization of *Staphylococcus pseudintermedius* associated with post-operative surgical infections in dogs. J Med Microbiol. 2015;64(9):1074–81 DOI:10.1099/jmm.0.000110.

5 Diribe O, North S, Sawyer J, Roberts L, Fitzpatrick N, La Ragione R. Design and application of a loop-mediated isothermal amplification assay for the rapid detection of *Staphylococcus pseudintermedius*. J Vet Diagn Invest. 2014;26(1):42–8 DOI:10.1177/1040638713516758.

6 Rubin JE, Ball KR, Chirino-Trejo M. Antimicrobial susceptibility of *Staphylococcus aureus* and *Staphylococcus pseudintermedius* isolated from various animals. Can Vet J. 2011;52(2):153–7

7 Weese JS, Poma R, James F, Buenviaje G, Foster R, Slavic D. *Staphylococcus pseudintermedius* necrotizing fasciitis in a dog. Can Vet J. 2009;50(6):655–6

8 Hanselman BA, Kruth SA, Rousseau J, Weese JS. Coagulase positive staphylococcal colonization of humans and their household pets. Can Vet J. 2009;50(9):954–8

9 Boost MV, So SY, Perreten V. Low rate of methicillin-resistant coagulase-positive staphylococcal colonization of veterinary personnel in Hong Kong. Zoonoses Public Health. 2011;58(1):36–40 DOI:10.1111/j.1863-2378.2009.01286.x.

10 Carroll KC, Burnham CD, Westblade LF. From canines to humans: Clinical importance of *Staphylococcus pseudintermedius*. PLoS Pathog. 2021;17(12):e1009961 DOI:10.1371/journal.ppat.1009961.

11 Lhermie G, La Ragione RM, Weese JS, Olsen JE, Christensen JP, Guardabassi L. Indications for the use of highest priority critically important antimicrobials in the veterinary sector. J Antimicrob Chemother. 2020;75(7):1671–80 DOI:10.1093/jac/dkaa104.

12 McCarthy AJ, Harrison EM, Stanczak-Mrozek K, Leggett B, Waller A, Holmes MA, Lloyd DH, Lindsay JA, Loeffler A. Genomic insights into the rapid emergence and evolution of MDR in *Staphylococcus pseudintermedius*. J Antimicrob Chemother. 2015;70(4):997–1007 DOI:10.1093/jac/dku496.

13 Smith JT, Amador S, McGonagle CJ, Needle D, Gibson R, Andam CP. Population genomics of *Staphylococcus pseudintermedius* in companion animals in the United States. Commun Biol. 2020;3(1):282 DOI:10.1038/s42003-020-1009-y.

14 Brooks MR, Padilla-Velez L, Khan TA, Qureshi AA, Pieper JB, Maddox CW, Alam MT. Prophage-Mediated Disruption of Genetic Competence in *Staphylococcus pseudintermedius*. mSystems. 2020;5(1) e00684–19 DOI:10.1128/mSystems.00684-19.

15 Penna B, Silva MB, Botelho AMN, Ferreira FA, Ramundo MS, Silva-Carvalho MC, Rabello RF, Vieira-da-Motta O, Figueiredo AMS. Detection of the international lineage ST71 of methicillin-resistant *Staphylococcus pseudintermedius* in two cities in Rio de Janeiro State. Braz J Microbiol. 2022;53(4):2335–41 DOI:10.1007/s42770-022-00852-9.

16 Phumthanakorn N, Prapasarakul N. Investigating the ability of methicillin-resistant *Staphylococcus pseudintermedius* isolates from different sources to adhere to canine and human corneocytes. Can J Vet Res. 2019;83(3):231–4

17 Bergot M, Martins-Simoes P, Kilian H, Chatre P, Worthing KA, Norris JM, Madec JY, Laurent F, Haenni M. Evolution of the Population Structure of *Staphylococcus pseudintermedius* in France. Front Microbiol. 2018;9:3055 DOI:10.3389/fmicb.2018.03055.

18 Balachandran M, Bemis DA, Kania SA. Expression and function of protein A in *Staphylococcus pseudintermedius*. Virulence. 2018;9(1):390–401 DOI:10.1080/21505594.2017.1403710.

19 Bruce SA, Smith JT, Mydosh JL, Ball J, Needle DB, Gibson R, Andam CP. Accessory Genome Dynamics of Local and Global *Staphylococcus pseudintermedius* Populations. Front Microbiol. 2022;13:798175 DOI:10.3389/fmicb.2022.798175.

20 Bankevich A, Nurk S, Antipov D, Gurevich AA, Dvorkin M, Kulikov AS, Lesin VM, Nikolenko SI, Pham S, Prjibelski AD, Pyshkin AV, Sirotkin AV, Vyahhi N, Tesler G, Alekseyev MA, Pevzner PA. SPAdes: a new genome assembly algorithm and its applications to single-cell sequencing. J Comput Biol. 2012;19(5):455–77 10.1089/cmb.2012.0021.

21 Gurevich A, Saveliev V, Vyahhi N, Tesler G. QUAST: quality assessment tool for genome assemblies. Bioinformatics. 2013;29(8):1072–5 DOI:10.1093/bioinformatics/btt086.

22 Treangen TJ, Ondov BD, Koren S, Phillippy AM. The Harvest suite for rapid core-genome alignment and visualization of thousands of intraspecific microbial genomes. Genome Biol. 2014;15(11):524 DOI:10.1186/s13059-014-0524-x.

23 Silva M, Machado MP, Silva DN, Rossi M, Moran-Gilad J, Santos S, Ramirez M, Carrico JA. chewBBACA: A complete suite for gene-by-gene schema creation and strain identification. Microb Genom. 2018;4(3) e000166 DOI:10.1099/mgen.0.000166.

24 Hyatt D, Chen GL, Locascio PF, Land ML, Larimer FW, Hauser LJ. Prodigal: prokaryotic gene recognition and translation initiation site identification. BMC Bioinformatics. 2010;11:119 DOI:10.1186/1471-2105-11-119.

25 Zhou Z, Alikhan NF, Sergeant MJ, Luhmann N, Vaz C, Francisco AP, Carrico JA, Achtman M. GrapeTree: visualization of core genomic relationships among 100,000 bacterial pathogens. Genome Res. 2018;28(9):1395–404 DOI:10.1101/gr.232397.117.

26 Feldgarden M, Brover V, Gonzalez-Escalona N, Frye JG, Haendiges J, Haft DH, Hoffmann M, Pettengill JB, Prasad AB, Tillman GE, Tyson GH, Klimke W. AMRFinderPlus and the Reference Gene Catalog facilitate examination of the genomic links among antimicrobial resistance, stress response, and virulence. Sci Rep. 2021;11(1):12728 DOI:10.1038/s41598-021-91456-0.

27 Zankari E, Allesoe R, Joensen KG, Cavaco LM, Lund O, Aarestrup FM. PointFinder: a novel web tool for WGS-based detection of antimicrobial resistance associated with chromosomal point mutations in bacterial pathogens. J Antimicrob Chemother. 2017;72(10):2764–8 DOI:10.1093/jac/dkx217.

28 van der Graaf-van Bloois L, Wagenaar JA, Zomer AL. RFPlasmid: predicting plasmid sequences from short-read assembly data using machine learning. Microb Genom. 2021;7(11) e000683 DOI:10.1099/mgen.0.000683.

29 Seemann T. Prokka: rapid prokaryotic genome annotation. Bioinformatics. 2014;30(14):2068–9 DOI:10.1093/bioinformatics/btu153.

30 Page AJ, Cummins CA, Hunt M, Wong VK, Reuter S, Holden MT, Fookes M, Falush D, Keane JA, Parkhill J. Roary: rapid large-scale prokaryote pan genome analysis. Bioinformatics. 2015;31(22):3691–3 DOI:10.1093/bioinformatics/btv421.

31 Brynildsrud O, Bohlin J, Scheffer L, Eldholm V. Rapid scoring of genes in microbial pan-genome-wide association studies with Scoary. Genome Biol. 2016;17(1):238 DOI:10.1186/s13059-016-1108-8.

32 Wishart DS, Han S, Saha S, Oler E, Peters H, Grant JR, Stothard P, Gautam V. PHASTEST: faster than PHASTER, better than PHAST. Nucleic Acids Res. 2023;51(W1):W443–W50 DOI:10.1093/nar/gkad382.

33 Payne LJ, Meaden S, Mestre MR, Palmer C, Toro N, Fineran PC, Jackson SA. PADLOC: a web server for the identification of antiviral defence systems in microbial genomes. Nucleic Acids Res. 2022;50(W1):W541–W50 DOI:10.1093/nar/gkac400.

34 Jolley KA, Maiden MC. BIGSdb: Scalable analysis of bacterial genome variation at the population level. BMC Bioinformatics. 2010;11:595 DOI:10.1186/1471-2105-11-595.

35 Mageeney CM, Lau BY, Wagner JM, Hudson CM, Schoeniger JS, Krishnakumar R, Williams KP. New candidates for regulated gene integrity revealed through precise mapping of integrative genetic elements. Nucleic Acids Res. 2020;48(8):4052–65 DOI:10.1093/nar/gkaa156.

36 Xia G, Wolz C. Phages of *Staphylococcus aureus* and their impact on host evolution. Infect Genet Evol. 2014;21:593–601 DOI:10.1016/j.meegid.2013.04.022.

37 Wang Y, Zhao N, Jian Y, Liu Y, Zhao L, He L, Liu Q, Li M. The pro-inflammatory effect of Staphylokinase contributes to community-associated *Staphylococcus aureus* pneumonia. Commun Biol. 2022;5(1):618 DOI:10.1038/s42003-022-03571-x.

38 Rooijakkers SH, Ruyken M, van Roon J, van Kessel KP, van Strijp JA, van Wamel WJ. Early expression of SCIN and CHIPS drives instant immune evasion by *Staphylococcus aureus*. Cell Microbiol. 2006;8(8):1282–93 DOI:10.1111/j.1462-5822.2006.00709.x.

39 Goshorn SC, Schlievert PM. Bacteriophage association of streptococcal pyrogenic exotoxin type C. J Bacteriol. 1989;171(6):3068–73 DOI:10.1128/jb.171.6.3068-3073.1989.

40 Arnold JW, Koudelka GB. The Trojan Horse of the microbiological arms race: phage-encoded toxins as a defence against eukaryotic predators. Environ Microbiol. 2014;16(2):454–66 DOI:10.1111/1462-2920.12232.

41 Shaikh N, Tarr PI. *Escherichia coli* O157:H7 Shiga toxin-encoding bacteriophages: integrations, excisions, truncations, and evolutionary implications. J Bacteriol. 2003;185(12):3596–605 DOI:10.1128/JB.185.12.3596-3605.2003.

42 Carey JN, Mettert EL, Fishman-Engel DR, Roggiani M, Kiley PJ, Goulian M. Phage integration alters the respiratory strategy of its host. Elife. 2019;8 e49081 DOI:10.7554/eLife.49081.

43 Wagner PL, Waldor MK. Bacteriophage control of bacterial virulence. Infect Immun. 2002;70(8):3985–93 DOI:10.1128/IAI.70.8.3985-3993.2002.

44 Smoot LM, Smoot JC, Graham MR, Somerville GA, Sturdevant DE, Migliaccio CA, Sylva GL, Musser JM. Global differential gene expression in response to growth temperature alteration in group A *Streptococcus*. Proc Natl Acad Sci U S A. 2001;98(18):10416–21 DOI:10.1073/pnas.191267598.

45 Fortier LC, Sekulovic O. Importance of prophages to evolution and virulence of bacterial pathogens. Virulence. 2013;4(5):354–65 DOI:10.4161/viru.24498.

46 Christie GE, Dokland T. Pirates of the Caudovirales. Virology. 2012;434(2):210–21 DOI:10.1016/j.virol.2012.10.028.

47 John J, George S, Nori SRC, Nelson-Sathi S. Phylogenomic Analysis Reveals the Evolutionary Route of Resistant Genes in *Staphylococcus aureus*. Genome Biol Evol. 2019;11(10):2917–26 DOI:10.1093/gbe/evz213.

