## Supplementary Figures for "Global phylogenomic analysis of *Staphylococcus pseudintermedius* reveals genomic and prophage diversity in multi-drug resistant lineages"

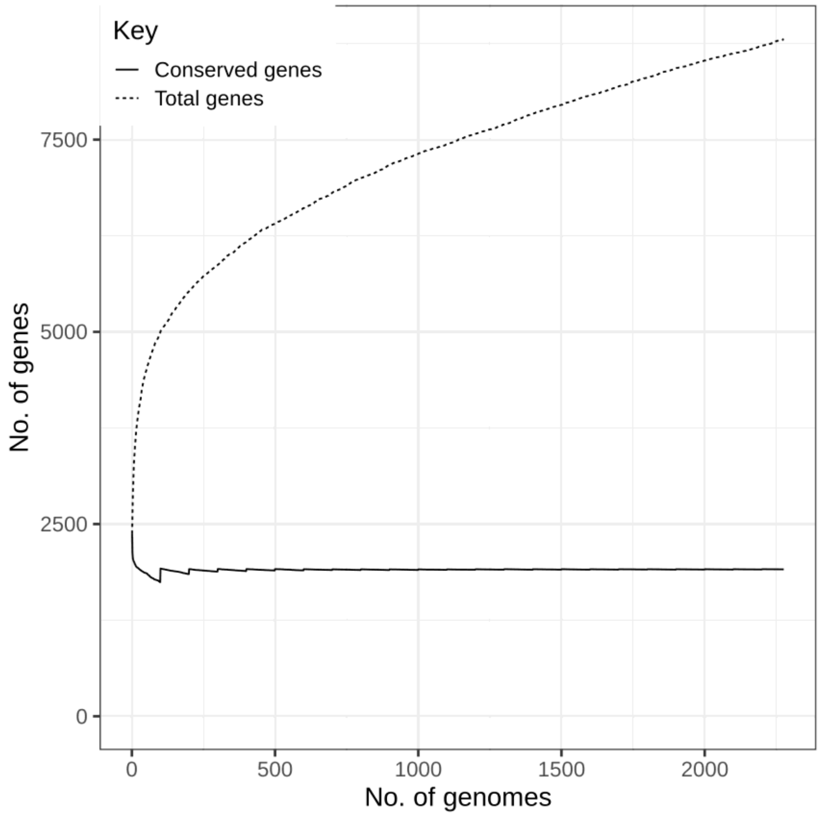


**Figure S1**: Gene accumulation curve. Analysis of the pangenome indicates the number of genes in the pangenome continues to increase with each additionally sequenced *S. pseudintermedius* genome included in the pangenome. Of the 6,814 accessory genes identified, 6,099 were only present 0-15% of genomes.

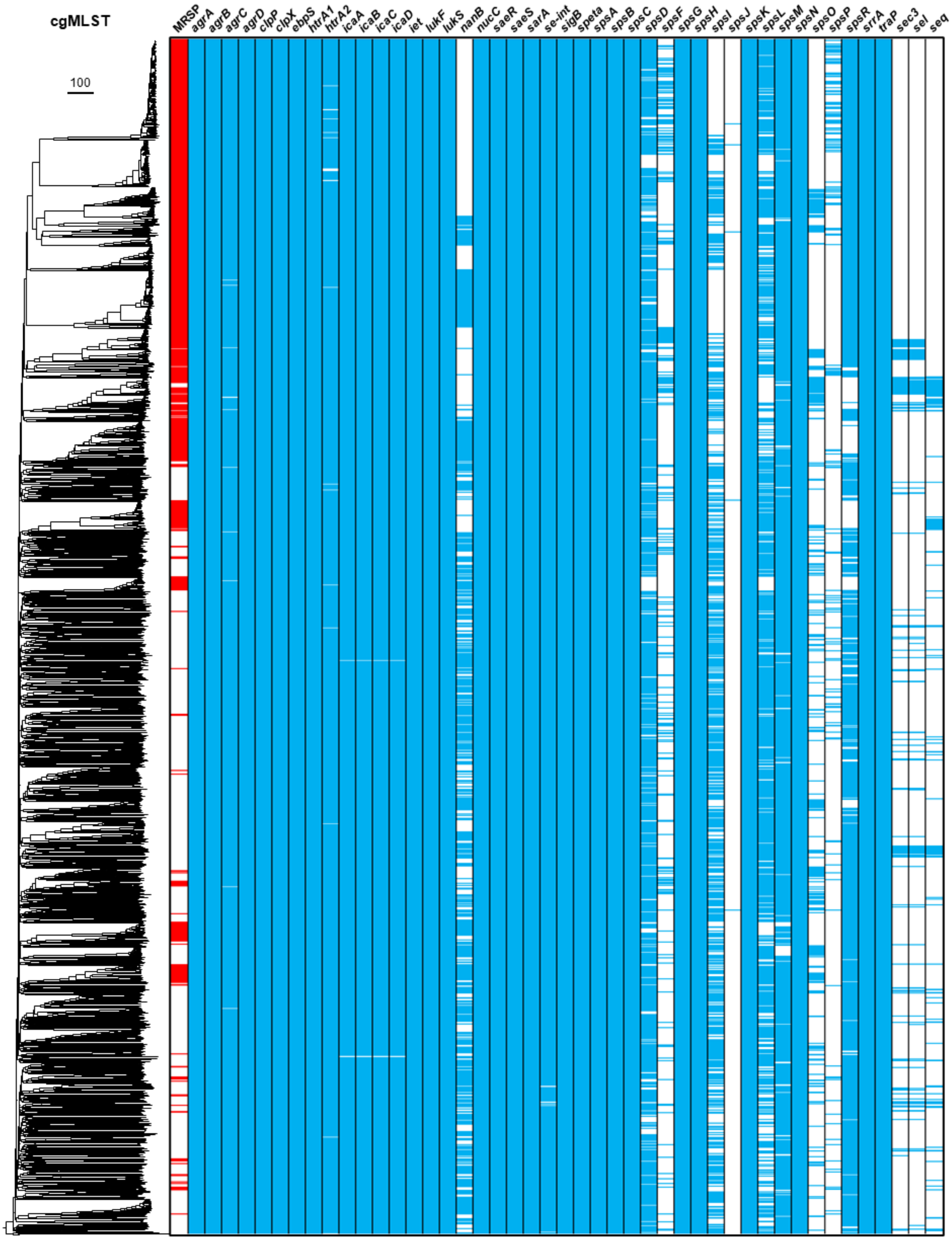


**Figure S2**: Maximum likelihood phylogeny showing the distribution of putative and reported virulence genes encoded by all 2,276 *S. pseudintermedius* genomes included in this study. Presence of a virulence gene is indicated by blue shading, absence is indicated by white shading. Full details of the phylogenetic distribution of virulence determinants are provided in Supplementary File 1.

Global phylogenomic analysis of Staphylococcus pseudintermedius reveals genomic and prophage diversity in multi-drug resistant lineages-Supplementary Material

A

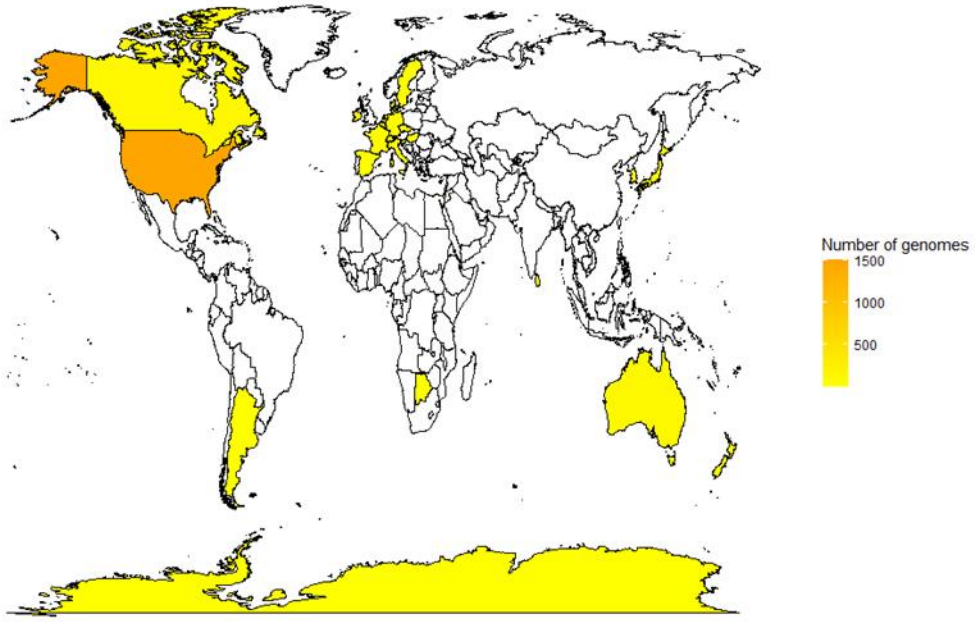


B

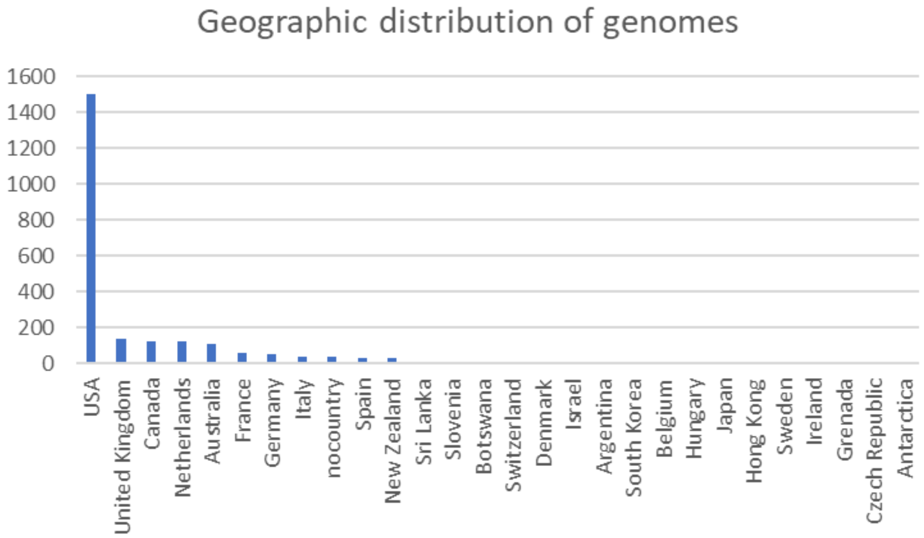


**Figure S3**: The geographic origin of isolates whose genome sequences were included in this study. **A**. Greater heat colour intensity indicates a greater number of genomes derived from that country. Most of the genomes included in this study are derived from USA. **B**. Bar chart showing the number of genomes derived from each country. Full details of the country of origin of each genome are provided in Supplementary File 1.

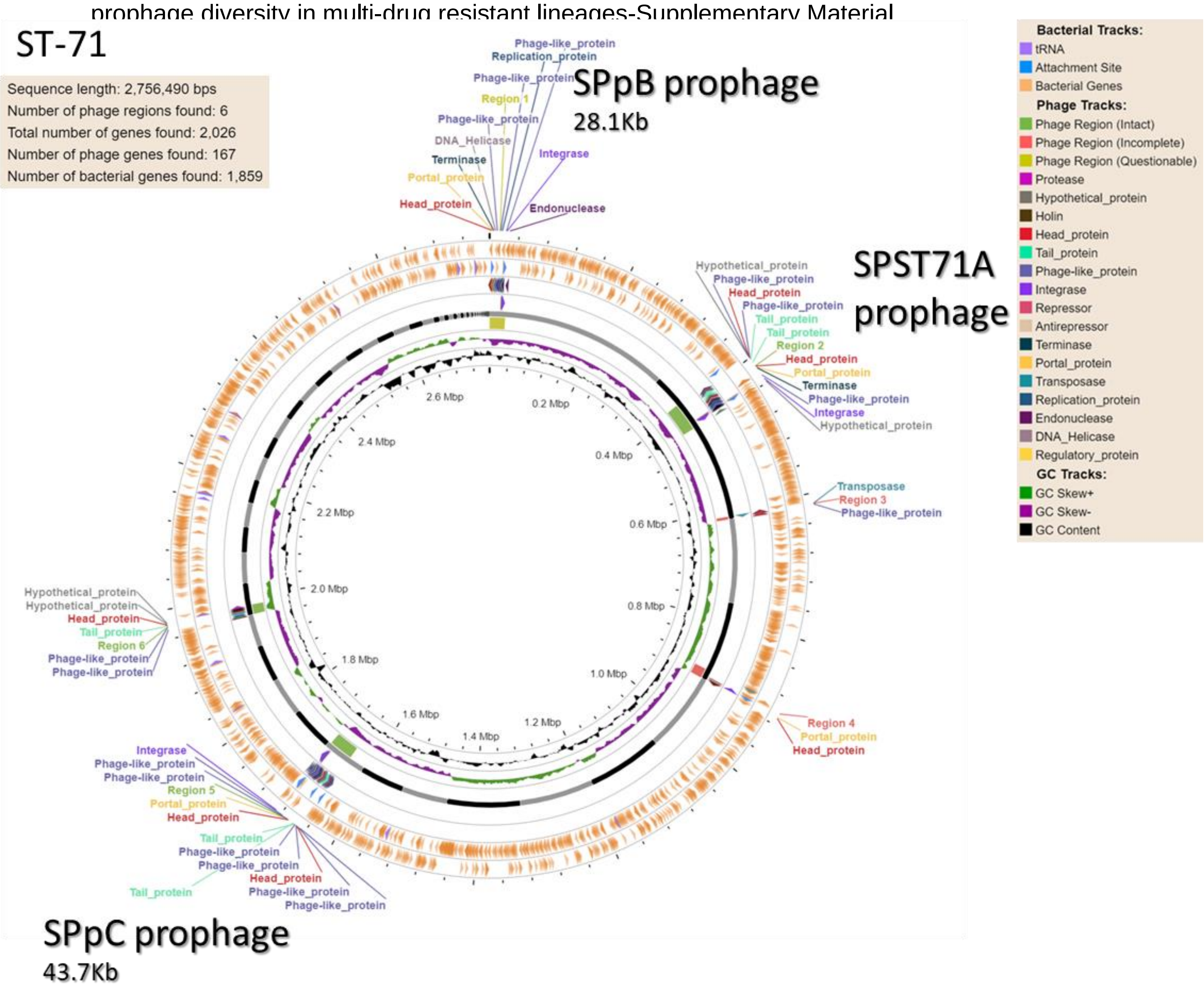


**Figure S4**: Circular genome map generated by PHASTEST, showing the number and location of intact prophages in a representative ST-71 genome (StaphpseudUoS10). Three large prophages SPST71A, SPpB, and SPpC have been identified in most ST-71 genomes which are absent from most other lineages.

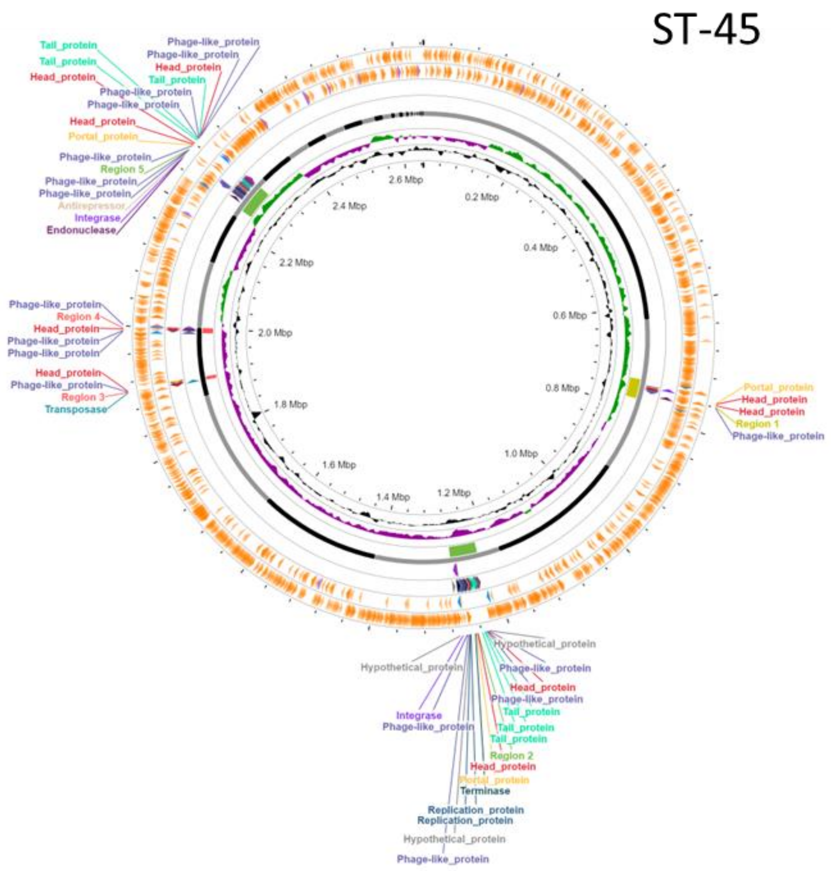


**Figure S5**: Circular genome map generated by PHASTEST, showing the number and location of intact prophages in a representative ST-45 genome. Most ST-45 genomes have at least 3 intact prophages, distinct from SPST71A, SPpB, and SPpC in ST-71.

Global phylogenomic analysis of Staphylococcus pseudintermedius reveals genomic and prophage diversity in multi-drug resistant lineages-Supplementary Material

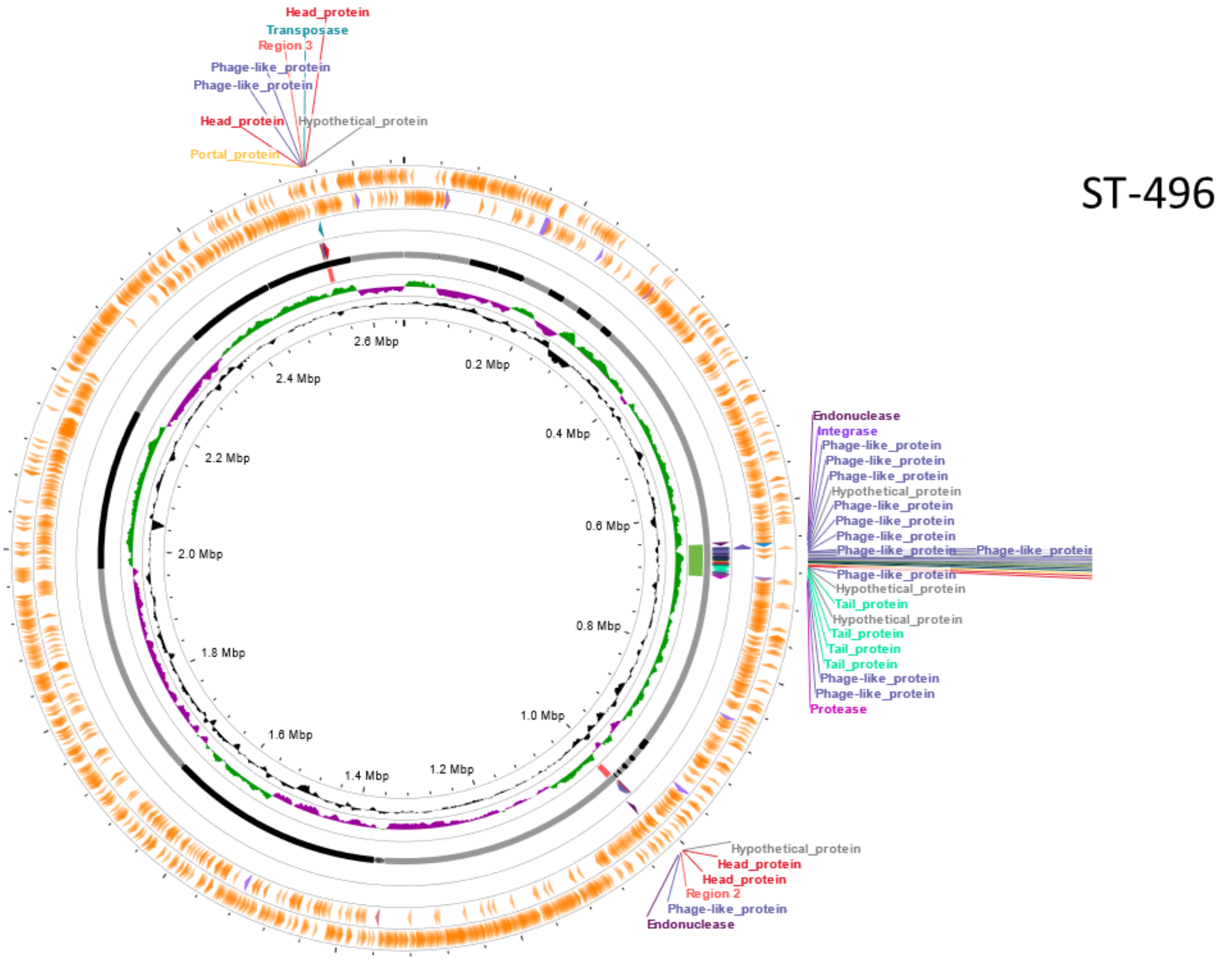


**Figure S6**: Circular genome map generated by PHASTEST, showing the number and location of intact prophages in a representative ST-496 genome. ST-496 only encode a single intact prophage.
